# A plastic attractor model of flexible rule-based selective attention

**DOI:** 10.1101/2025.09.07.674747

**Authors:** Christopher J. Whyte, Sanjay G. Manohar, Eva Feredoes, Alexandra Woolgar

## Abstract

A defining feature of human cognition is the ability to select and respond to stimuli flexibly in different circumstances. Here we show that a recently proposed class of small associative neural network, plastic attractors, can perform such flexible cognitive functions through the rapid formation of task-based attractors. We simulated a rule-based selective attention paradigm, in which agents respond to one task-relevant feature of a visual stimulus, while ignoring another, irrelevant feature, and respond to the attended feature according to a predefined stimulus-response mapping rule. The model consists of a broadly tuned prefrontal population with rapidly changing recurrent connections to sensory neurons that compete via lateral inhibition. In this framework, the rules governing the focus of selective attention are not coded explicitly, but arise as an emergent property of temporary associations between stimulus features and motor responses. The model exhibited activation properties that embody cornerstone concepts in current attentional theory including mixed selectivity, adaptive coding and biased competition, and reproduced a number of classic behavioral and neural findings. A causal test of the model using non-invasive brain stimulation concurrent with functional magnetic resonance imaging (TMS-fMRI) in humans showed that network perturbation reproduced neural decoding and behavioural data. When features were task-relevant, they rapidly formed synaptic connections with frontal cortex binding them into an attracting state, which manifest as prioritized representation of attended information, but this state was readily corrupted by brain stimulation. The model shows mechanistically that rapid synaptic changes could explain flexible rule-based control of attention.

## Introduction

Flexible selective attention is a foundational feature of human cognition. Based upon the rules of our current task we can prioritise the processing of stimulus information that is relevant to the task at hand, ignore irrelevant information, and use the relevant information to guide action selection, all whilst retaining the capacity to rapidly switch tasks. This process is thought to depend on a frontoparietal cognitive control or “multiple demand” network (Cole and Schneider 2007, Duncan 2010, Gratton et al. 2018, Assem et al. 2020, Soltani and Koechlin 2022) which adaptively codes task relevant information (Duncan 2001, Zheng et al. 2024) and integrates it into a task specific structure (Duncan et al. 2020, Ito et al. 2022, Cole 2024, Duncan 2024), and which, in turn, provides feedback to sensory cortices, biasing competition among stimulus elements in favour of attended information (Desimone and Duncan 1995).

In line with this broad hypothesis, work in both human neuroimaging and non-human primate electrophysiology has shown that across sensory and frontoparietal cortex the neuronal representation of attended (i.e. task relevant) stimuli (or stimulus features) is prioritised at the expense of unattended stimuli in terms of both neural firing rates (e.g., Moran and Desimone 1985, Chelazzi et al. 1993, Desimone and Duncan 1995, Luck et al. 1997, Chelazzi et al. 1998) and multivariate measures of information coding (e.g., Jehee et al. 2011, Gratton et al. 2013, Woolgar et al. 2015, Jackson et al. 2017, Jackson and Woolgar 2018, Chen et al. 2021, Moerel et al. 2024, Zheng et al. 2024, Dermody et al. 2025). Neural codes in frontoparietal cortex, in particular, appear to be highly flexible. For example, in non-human primates, single neurons have long been known to adjust their selectivity profiles to reflect task-relevent stimulus boundries of the momentarily relevent stimulus (Rao et al. 1997, Duncan 2001, Freedman et al. 2001). This effect is also seen in human neuroimaging. For example, activity in frontoparietal control regions appears to shift to prioritise currently relevant stimulus distinctions (e.g., Jackson et al. 2017, Dermody et al. 2025), or to emphasise different task aspects based on difficulty (e.g., Woolgar et al. 2015, Wisniewski et al. 2023), with single voxels in these regions contributing to multiple classification dimensions (e.g., Jackson and Woolgar 2018, Jackson et al. 2025) (for reviews see Woolgar et al. 2016, Zheng et al. 2024). This dynamic selectivity is mirrored by flexible connectivity between frontoparietal and other brain systems (Cole et al. 2013, Cole 2024), with brain wide connectivity profiles predicting information content in frontoparietal cortex (Schultz et al. 2022).

In addition, there is converging evidence from both human and non-human primates that activity in frontoparietal regions influences attention-depedent stimulus coding in sensory cortex. For example, in attention tasks, Granger-causal analysis of multivariate codes show top-down influence of PFC codes on the structure of representations in visual cortices (Snyder et al. 2021, Goddard et al. 2022). In non-human work, it has long been observed that stimulating frontal regions such as the frontal eye fields modulates stimulus-driven activity in sensory cortices (Moore and Armstrong 2003). Similarly, modulating human prefrontal activity with transcranial magnetic stimulation (TMS), modulates activity (Ruff et al. 2006), excitability (Veniero et al. 2021) and multivariate coding of attended stimulus information (Jackson et al. 2021) in visual cortices.

Finally, there is both behavioural and neural evidence that the timescale over which these codes are assembed and adjusted is fast, on the order of a single trial. A single exposure to task instructions allows human participants to immediately excute decision making tasks (Cole et al. 2013), and neural activity in non-human primate lateral PFC is adjusted on a single-trial timescale along with the formation and distruction of new associations (Bocincova et al. 2022). Similarly, in human neuroimaging, attentional prioritisation occurs within the first 200ms of stimulus onset (e.g., Battistoni et al. 2020, Grootswagers et al. 2021, Goddard et al. 2022, Moerel et al. 2022, Lu et al. 2025), before decision making can begin (Moerel et al. 2024), even if attention is switched mid-trial in a multi-step task (Barnes et al. 2021).

Despite these well-characterised neural properties, computational models that explain *how* these signatures of flexibility arise are lacking. Feedforward networks and recurrent networks with slow changing or fixed weights allow task-selective units to direct attention to relevant features to drive decisions (e.g., Cohen et al. 1990, Dehaene and Changeux 1991, Botvinick and Cohen 2014, Denison et al. 2021). However they typically either need hard-coded task neurons with fixed neural selectivity, rely on biologically-implausible gradient based learning schemes (e.g., Flesch et al. 2022), or require training periods that take place over timescales much longer than the single trial exposure to task instructions required to account for human performance (e.g., OʹReilly and Frank 2006), or do not aim to model the task acquisition process (e.g., Denison et al. 2021).

A parallel literature heavily implicates frontoparietal cortex in the learning, maintenance, execution, and switching of task rules, including stimulus-response rules that define the behavioural actions needed for each stimulus (e.g., Bunge et al. 2003, Wallis and Miller 2003, Badre and DʹEsposito 2007, Koechlin and Summerfield 2007, Bode and Haynes 2009, Woolgar et al. 2015, Woolgar et al. 2016, Gonzalez-Garcia et al. 2017, Palenciano et al. 2019, Gonzalez-Garcia et al. 2021). Although stimulus-response rule application and selective attention are often considered in separate literatures, or are combined with an emphasis on the effect of attention on rule use (e.g., Gonzalez-Garcia et al. 2020), a parsimonious possibility is that, in many circumstances, selective attention – the prioritisation of task relevant aspects of stimuli – could arise directly from the process of forming appropriate stimulus-response associations. This is the possibility that we address in this paper.

Through numerical simulations of a recently proposed class of associative recurrent neural network, *plastic attractors*, we show how a flexible population of prefrontal-like ‘conjunction’ neurons with winner-take-all dynamics that share recurrent connections with a population of sensory-like ‘feature’ neurons, can rapidly learn, and perform, rule-based selective attention tasks through fast Hebbian plasticity. Specifically, the presentation of task instructions, consisting of a stimulus-response mapping, is rapidly encoded in a fixed-point attractor through the rapid modification of synaptic weights, maintained in persistent activity, and then stored in an ‘activity silent’ manner where it can be rapidly (and reliably) recalled by an appropriate cue to guide subsequent neural activity. This combination of rapid Hebbian plasticity, persistent activity, and activity silent storage has been proposed as a potential mechanism of working memory (Howard and Kahana 2002, Sandberg et al. 2003, Fiebig and Lansner 2017). Indeed, Manohar et al. (2019) showed that the plastic attractor architecture could account for nearly 30 established findings in the working memory literature, with concomitant simulation of neural decoding and stimulation.

Here we show that, with only minor modification, the plastic attractor architecture generalises to rule-based decision-making tasks, and in doing so, gives rise to canonical selective attention effects. To establish the validity of the model we simulate a well-studied rule based perceptual decision making task and confirm that the model reproduces the key qualitative features of empirical behavioural and neural (fMRI and MEG) data from human participants including cross-block congruency effects in reaction-time, and the prioritisation of attended (i.e. task relevant) information in both firing rates and multivariate decoding. We then provide a novel causal test of the model by simulating a unique concurrent TMS-fMRI experiment (Jackson et al. 2021) in which activity in dorsolateral prefrontal cortex (dlPFC) was perturbed whilst participants performed a rule-based perceptual decision making task in the fMRI scanner. We found that transient perturbation of the model’s prefrontal-like conjunction neurons reproduced the key neural effects of TMS of reducing decoding accuracy for task relevant, but not task irrelevant, information. Moreover, the model provided a new lens to resolve the apparently paradoxical relationship between the reported neural and behavioural effects, with the effect of stimulus congruency on reaction times (RTs) reduced, after TMS, due to weakened attractor states. The data suggest that the rapid synaptic changes that underpin the operation of the plastic attractor could underlie flexible prefrontal control of attention.

## Methods

### Model Dynamics

The network employed in this paper is a (slightly) modified version of the plastic attractor network introduced by Manohar et al. (2019). The evolution of the network is described by the following difference equations:

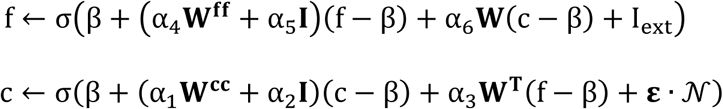

Where c and f denote conjunction and feature unit activation vectors whose values are bounded by σ(·) which is a piecewise function that saturates at values of 0 and 1 and is linear otherwise:

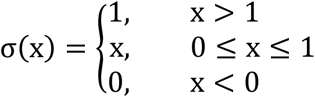

β is the baseline neuronal activity vector and is set to .175, and **W**^**cc**^and **W**^**ff**^are the conjunction to conjunction, and feature to feature synaptic weight matrices. **W**^**cc**^is a matrix of 1’s, and the entries of **W**^**ff**^are equal to 1 if they are in the same feature dimension, and 0 otherwise. **W** is the feature to conjunction weight matrix, and its transpose is the conjunction to feature weights. **W** is initialised with random values drawn from a uniform distribution over the interval [0,1]. Each of the weight matrices were scaled by the following six free parameters; α_1_and α_4_ - lateral inhibition, α_2_and α_S_ - self-excitation, and α_3_ and α_6_ - synaptic gain, with values α = [-0.45, 1.0, 0.08, -0.28, 0.73, 0.04]. The sign of the parameters were chosen so that the diagonal entries of the conjunction-to-conjunction, and feature-to-feature weight matrices were positive, and off diagonal elements were negative. This caused lateral inhibition between units when the firing rates were above the baseline value β. This specific selection of parameter values was adapted from the ‘high accuracy regime’ of Manohar et al, (2019) with two minor modifications. First, we reduced the value of the lateral inhibition parameter for the conjunction units from -0.5 to -0.45, so that the simulated TMS pulses (described below and in simulation 2) did not result in seizure-like behaviour. Second, we slightly increased the self-excitation parameter for feature units from 0.7 to 0.73, so that the network was able to recover after receiving a simulated TMS pulse. Finally, *N* is a Gaussian noise term scaled by the parameter ε (which was set to 0.005 following Manohar et al., 2019), and I_ext_ is the external input vector the values of which we kept between -1 and 1.

The weight matrix **W** connecting feature and conjunction units was updated according to the following Hebbian covariance rule:

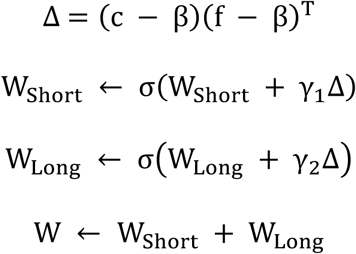

Where Δ is the Hebbian update, and **W** is the sum of two weight matrices **W**_Short_ and **W**_Long_. **W**_Short_ is updated with a learning rate of γ_1_ = 0.02, and is bound to values on the interval [0,1]. In contrast, **W**_Long_is updated with a slower learning rate of γ_1_= 0.0002, and is bound to the much narrower interval [0,0.2]. The addition of the long term weights is the only major deviation from the model structure presented by (Manohar et al., 2019). The long term weight matrix acted as a bias and helped keep the selectivity of the conjunction units stable across blocks once they came to encode a particular set of features (**Figure S1**). Indeed, without the addition of the long term weights the model was not able to capture the behavioural congruency effect (**Figure S1**), which in the tasks we aimed to simulate, required memory traces to be carried across blocks. Secondly, adding long term plasticity also prevented the conjunction units changing their selectivity profiles over blocks. Both of these features align well with empirical results. Specifically, behavioural congruency effects are well established, including in the tasks we aimed to simulate (Jackson et al. 2021), and at the population level, selectivity – mixed or otherwise – must be somewhat stable over time, as fMRI decoding (including from the frontoparietal regions) cross-validates across runs. To check that the addition of the second slower time-scale of plasticity did not interfere with the capacity of the model to perform the original working memory task it had been developed for, we verified that it could still perform the task used throughout Manohar et al, (2019), maintaining the general validity of the model (**Figure S2**).

## Simulation Protocol and Tasks

### Simulation of rule-based selective attention task

We used the model to simulate a classic rule-based feature-selective (sometimes called feature-specific) attention paradigm in which participants make perceptual decisions about separate features of single objects (Garner 1978, Rossi and Paradiso 1995, Nobre et al. 2006). These paradigms have been used extensively to study the neural basis for rule-based selective attention (Anllo-Vento and Hillyard 1996, Kauramäki et al. 2007, Altmann et al. 2008, Chen et al. 2012, Downer et al. 2017, Jackson et al. 2017, Jackson et al. 2021, Goddard et al. 2022, Dermody et al. 2025, Lu et al. 2025) and reflect the ubiquitous cognitive requirement of selecting one relevant feature of complex stimuli while ignoring others. The simulated tasks had the model make a perceptual decision about one of two orthogonal stimulus feature dimensions, nominally, colour (green and blue), and shape (square, circle), according to the rule presented on each block. In half of the blocks, the network was instructed to generate a response based upon the colour of the stimulus, and in the other half, it was instructed to generate a response based on the shape of the stimulus. The rule consisted of a mapping between the relevant feature and a motor response.

To simulate this task we used a network with four conjunction units, and six feature units two of which were labelled as motor responses (**Figure 1**). Each block began by presenting the network with a set of task rules by coactivating each stimulus feature unit in the relevant dimension and the appropriate motor response (i_ext_ = + 1), deactivating the non-active stimulus feature in the relevant dimension and the inappropriate motor response (i_ext_ = - 1), and partially activating both features of the irrelevant dimension (i_ext_ = + 0.5). Each rule consisted of two stimulus-response mappings each of which was presented separately. For example, for the rule segment “press left for green”, we used the input vector I_ext_ = [+1, -1, +0.5, +0.5, +1, -1] where the first two elements correspond to colours (blue, green), the third and fourth to shapes (square, circle), and the fifth and sixth to motor responses (left, right). This mimicked the visual instruction to press left for green objects of either shape as presented to humans in Jackson et al. (2021). Each of the two rule segments (e.g., “press left for green” and “press right for blue”) was presented for 250 time-steps, followed by an interstimulus interval of 150 time-steps in which all of the feature units were deactivated (i_ext_ = - 1). During the stimulus period, the model was presented with twelve trials, consisting of 3 presentations of each of the four stimulus combinations (i.e. green square, green circle, blue square, blue circle) in pseudorandom order. Stimulus presentation on each trial consisted of maximally activating the relevant stimulus feature units (i_ext_ = + 1). This was followed by a response period where all input into the feature units was removed (i_ext_ = 0), and the network was allowed to generate a response via pattern completion. Finally, in the interstimulus interval all the feature units were deactivated (i_ext_ = - 1) returning the network to baseline. Each stimulus was presented for 50 time-steps, followed by a response period of 250 time-steps and an interstimulus interval of 100 time-steps. We simulated 20 blocks of 12 trials with 20 random seeds.

**Figure 1.**
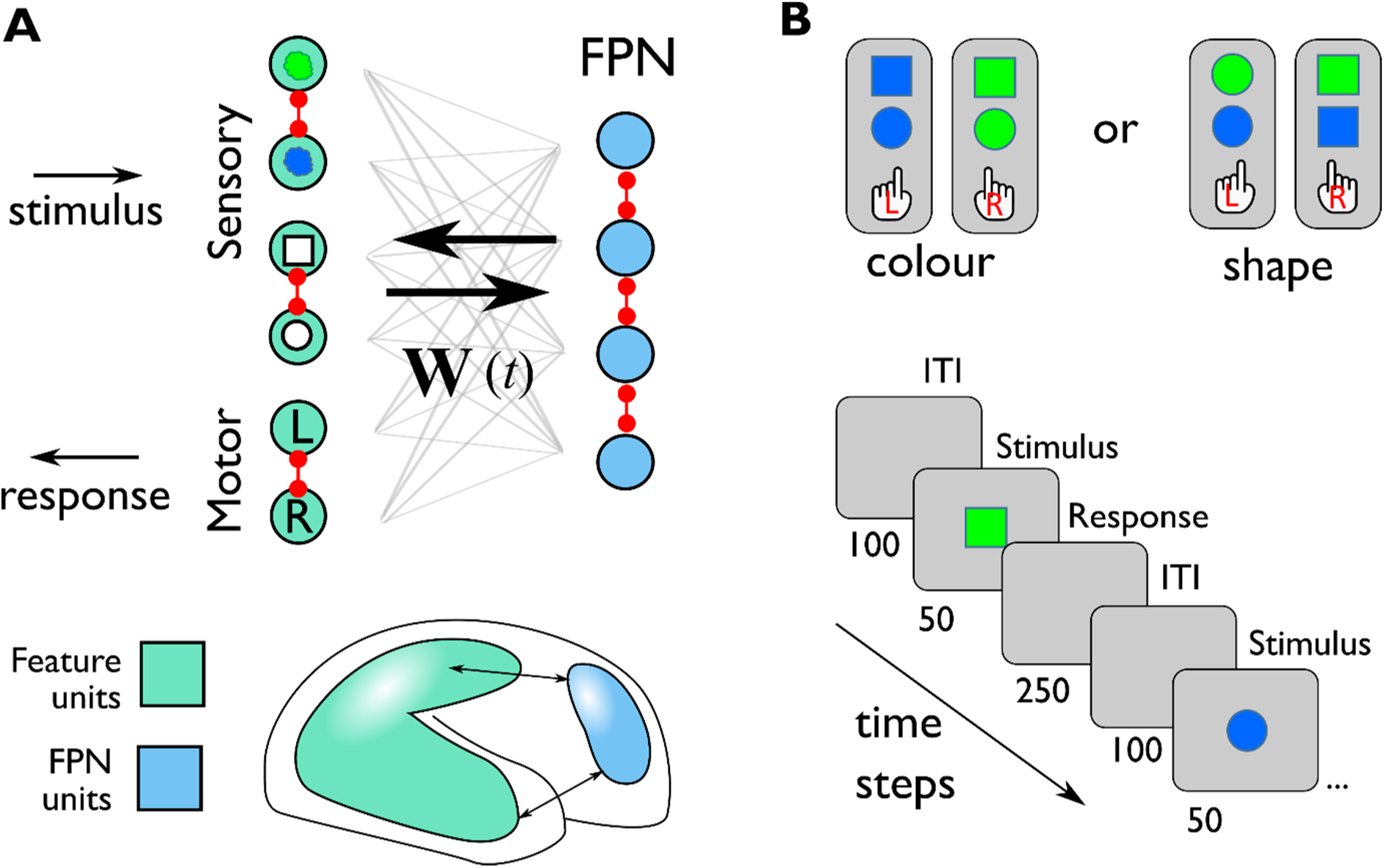
Model architecture and simulated task. **A** Architecture of the plastic attractor network consisting of two varieties of neurons, feature units (teal), which have fixed selectivity and correspond to sensory, conceptual, or motor representations in sensorimotor cortex, and conjunction units (blue), which have broad all-to-all bidirectional connectivity, initialised with random synaptic weights that, through the action of the Hebbian learning rule, quickly come to be selective for a subset or “conjunction” of feature units. Red connections signify lateral inhibition: Feature units compete within feature dimensions (e.g. colour, shape, response), and conjunction units have blanket competition leading to winner-take-all dynamics. **B** Feature-based attention task. Rules consisting of the relevant mappings between stimulus features (green, blue, square, circle) and responses (left, right) were sequentially presented to the network at the beginning of each block. Each rule dictates which dimension (colour or shape) is task-relevant (i.e. should be attended to) and which dimension is task-irrelevent (i.e. should be ignored). The rule presentation period was followed by twelve trials consisting of four psuedorandomised presentations of each stimulus combination (green-circle, green-square, blue-circle, blue-square).

### Simulation of TMS

In the second simulation we used the same task, but we now added a simulation of direct neural stimulation of the conjunction units by TMS. This was done to simulate the paradigm of Jackson et al. (2021) who used concurrent TMS-fMRI, a non-invasive technique allowing for the simultaneous perturbation and measurement of human neural activity (e.g., Bergmann et al. 2021, Woolgar et al. 2025), to examine the causal role of right dlPFC in driving feature-selective attention. In that study, the authors delivered a series of TMS pulses to the dlPFC during stimulus presentation in a task analogous to the one described above. Here, we did not aim to simulate the biophysical effects of TMS, which are unlikely to be captured by the network of rate neurons. Instead, we aimed to reproduce the phenomenology of TMS which includes a brief burst of excitation followed by a prolonged period of inhibition (e.g., Romero et al. 2019), in order to observe its effects on the model. We achieved this by maximally activating all conjunction units during the stimulus presentation period. Like the experimental data, after the brief period of maximal activation, the lateral inhibition between conjunction units caused all conjunction unit activity to drop and the conjunction units to re-compete for dominance. This had the effect of transiently reducing feedback from conjunction to feature units disrupting the recurrent circuit underlying maintenance and response generation. Because we did not have an *a priori* hypothesis about the correspondence between the parameters of the TMS protocol used by Jackson et al. (2021) and the simulation parameters, we ran the model over a range of TMS durations, referred to as the TMS “dose”, which can be thought of an analogous to a range of TMS stimulation intensities. We held the activation of these units for between 1 to 19 timesteps ending at stimulus offset. For example, a pulse of length 1 turned on at the final time-step of stimulus presentation, whilst a pulse length of 19 turned on 19 time-steps before stimulus offset. For the main analysis we applied TMS on every trial. However, we also examined the effect of interleaving TMS with non-TMS trials, which gave qualitatively similar results (**Figure S3**). In all other respects, the task of simulation 2 was identical to simulation 1.

The simulations were written in Python based upon the equations and parameters supplied in the supplementary material of Manohar et al. (2019).

## Model Analysis

### Model accuracy and RT

Following Manohar et al. (2019), we defined the network’s response as the maximally activated motor response during the response period, and the reaction time (RT) as the time to 98% of the maximum activation of that unit occurring more than 10 timesteps after the stimulus offset (i.e., during the response period). The model’s behavioural choice was the motor feature unit that was active at this timepoint, and was deemed correct if it matched the correct response that should have been generated based on the stimulus and currently active rule.

### Information in model units

To compare the simulated neural activity generated by the model to results from human neuroimaging, we applied a multivariate decoding pipeline similar to that used in the analysis of magnetoencephalography (MEG) and electroencephalography (EEG) (e.g., Grootswagers et al. 2017). Specifically, we used logistic regression with 5-fold cross-validation (unless noted otherwise) to decode stimulus features (i.e. green vs blue and circle vs square) within each dimension (shape and colour) from both conjunction and feature unit activity. The weights and intercept were fit using the L-BFGS solver in the sklearn.linear_model package (version 1.0.2) for Python 3 (Pedregosa et al. 2011).

### Quantification of synaptic effects

To quantify changes in the synaptic weights as a consequence of TMS, we reasoned that in networks consisting of simple linear dynamical systems the attractors of a system are described by the eigenvalues of the weight matrix (which describe the asymptotic behaviour of the system). However, our network is both non-linear and non-square (there are four conjunction units and six feature units). To study the effect of simulated TMS on the weight matrix we, therefore, approximated the full model with a reduced set of coupled difference equations. Specifically, we removed the non-linearity, lateral inhibition, input, noise, and additive constants, leaving us with:

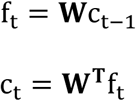

Importantly this reduction permits one further simplification, as c_t-1_ =

**W**^**T**^f_t-1_, allowing us to write the network in terms of a single difference equation that we can understand by studying the eigenvalues of the (now square) weight matrix

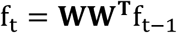

The behaviour of difference equations such as this can be described through a diagonalisation of the matrix **WW**^**T**^. Specifically, f_t_ = **X**Λ^**t**^**X**^-1^f_O_. Where **X** is a matrix containing the eigenvectors of **WW**^**T**^, and Λ is a matrix of 0’s containing the eigenvalues of **WW**^**T**^ down the diagonal. As t → ∞, states corresponding to eigenvalues greater than 1 blow up, eigenvalues equal to 1 correspond to steady states, and eigenvalues less than 1 decay to 0. Studying the weight matrix of this highly reduced network, therefore, tells us what feature units are amplified by the recurrent circuit connecting conjunction and feature units. In the context of the full network, the values of f_t_ are bounded between 0 and 1, and lateral inhibition punishes the non-winning stimulus elements within each dimension. The attractors, therefore, correspond to the patterns of features that are amplified by the weight matrix. Since the feature vectors corresponding to each stimulus-response mapping under each rule are linearly independent, we know that to complete the task the weight matrix

**WW**^**T**^needs to have at least two amplifying eigenvalues (i.e. two eigenvalues greater than 1). To measure this in the network we recorded the number of eigenvalues of the matrix **WW**^**T**^that were > 1 at the end of each block of trials.

## Results

### Face validity of the model

#### The model performs the task with high accuracy, and response times modulated by stimulus-response congruency

To establish the face validity of the model we first simulated a simplified version of the rule-based feature-selective attention task that we had used previously in human neuroimaging experiments using MEG (Barnes et al. 2021, Goddard et al. 2022) and fMRI (Jackson et al. 2017, Jackson et al. 2021, Dermody et al. 2025). As described in the Methods, each block started by presenting the task-rules to the model. Each rule dictated which of the two stimulus dimensions, colour or shape, was relevant, and how each feature along this dimension (e.g., blue, green) should be associated with each response (left, right). Following the presentation of the rules, the model was presented with twelve trials consisting of three instances of each of the four stimulus combinations in pseudorandom order. The model performed the task at near ceiling accuracy (M = 96.4%, SEM = 0.003), with an average RT of 39.8 time-steps for correct trials (SEM = 0.4). As for human participants (Jackson et al. 2021), the model also showed an effect of congruency, with shorter RTs for stimuli where both the relevant and irrelevant dimension mapped to the same motor response (green square and blue circle) compared to stimuli where the relevant and irrelevant features mapped to different motor actions (green circle and blue square); average congruent RT = 30.14 time-steps (SEM = 0.48), average incongruent RT = 50.19 time-steps (SEM = 0.94).

### Task relevant information is prioritised in silico, as in human data

Inspection of the model activity (**Figure**) showed that under each of the two rules, activity in the two feature units corresponding to the input was matched during the stimulus presentation period (seen as black bars in Feature Units heatmap). But, after the removal of the stimulus, activity was greater for the stimulated unit in the relevant stimulus dimension (e.g., the unit for “green”) than the unit in the irrelevant stimulus dimension (e.g., the unit for “circle”). Feature neurons coding for the presented stimulus on the relevant dimension retained a high level of activity even after the stimulus was removed (e.g., “green”/“blue” in **Figure 2A**). By contrast, in the irrelevant dimension activity was lower, and roughly equivalent between the presented and non-presented stimulus (e.g., “circle” and “square” in **Figure 2A**).

**Figure 2.**
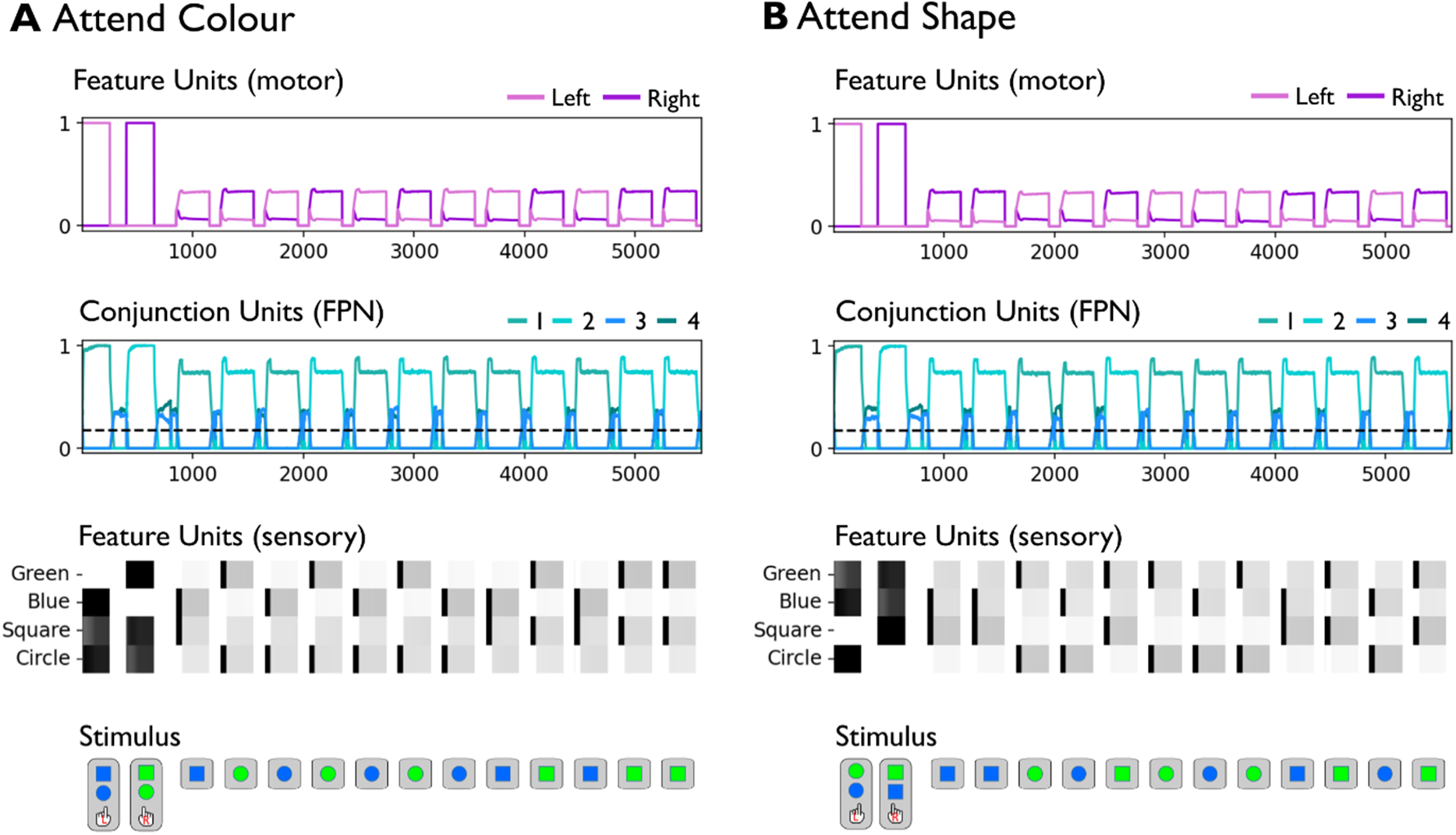
Model activity for the attend colour and attend shape tasks. **A** Model activity for the attend colour task. Simulated input (stimulus) is shown on the lowest row. Above this, raster-style plots show activity in the sensory feature units. Middle line plots conjunction unit activity, and top line plots show motor responses. Although motor units are modelled as a variety of feature unit, we separated them in the presentation here for ease of interpretability. Each colour denotes a distinct conjunction and feature unit. The first two epochs in the figure show the presentation of the two stimulus response mappings under each rule. For example, the first epoch associates the blue colour with the left response (pink), which come to be associated by the #1 conjunctive unit (light green). **B** Model activity for the attend shape rule, conventions as in A.

We trained a linear classifier to distinguish between each feature within the two stimulus dimensions (i.e. green vs blue, square vs circle) for both conjunction and feature units (**Figure 3**). We carried this out separately when the stimulus dimension was relevant (e.g., colour in the attend colour task) and when it was irrelevant (e.g., colour in the attend shape task) and averaged the resulting classification scores over shape and colour to give a single decoding accuracy for relevant and irrelevant features at each timepoint. Decoding accuracy for the simulated data displayed the same key qualitative characteristics as the empirical data. For conjunction units, decoding accuracy was higher for relevant than for irrelevant information throughout the trial (**Figure 3A**). This matched the pattern observed, using a similar decoding pipeline and task (Goddard et al. 2022), in human MEG data arising from the frontal lobe (**Figure 3B**). In the model’s feature units, decoding accuracy was high and similar for relevant and irrelevant information during stimulus presentation, but following the removal of the stimulus, only decoding of relevant information was sustained whilst decoding of irrelevant information dropped substantially (**Figure 3C**). This closely matched the pattern observed in the posterior ROI (over occipital/visual cortex) of the human MEG data, where decoding accuracy for relevant and irrelevant stimuli was matched during the first feedforward sweep, but only task-relevant information displayed above chance decoding for a sustained period (**Figure 3D**). Thus the model matched both the timecourse and spatial arrangement of human neural data. The simulated data also provided a qualitative match to a number of fMRI studies reporting stronger coding of relevant relative to irrelevant information in frontoparietal control and visual brain regions e.g., (Jackson et al. 2017, Jackson and Woolgar 2018, Chen et al. 2021, Dermody et al. 2025). In particular, these studies do not provide temporal resolution, but report that decoding accuracy for relevant information is often overall stronger than coding of irrelevant information in both neural systems, and as in the model, and irrelevant information tends to be at or only slightly above chance in the frontoparietal multiple demand regions, as we observed in the model’s conjunction units.

**Figure 3.**
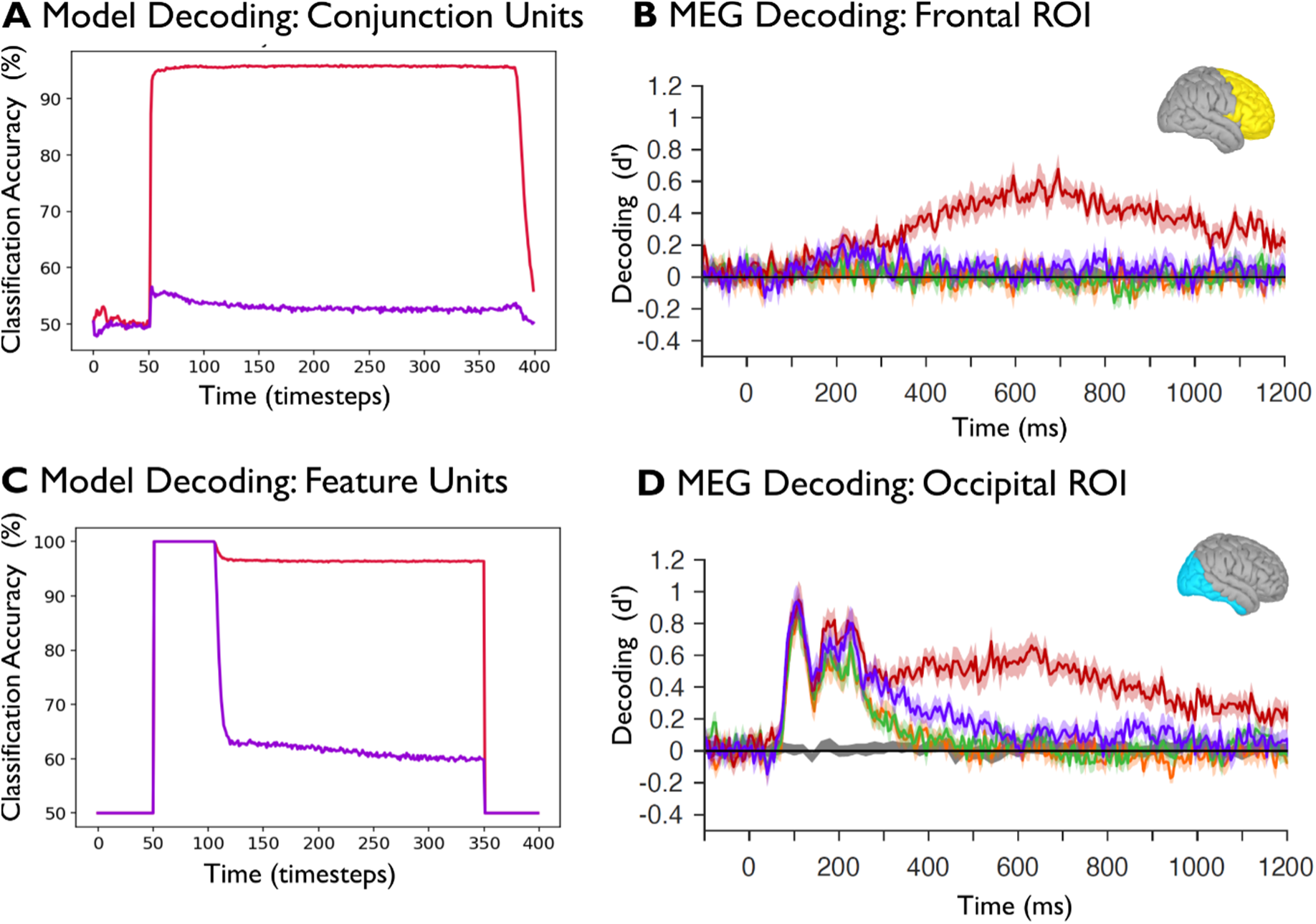
Decoding of relevant and irrelevant information. **A** Stimulus decoding accuracy from conjunction units for the relevant (red, e.g., shape during shape task) and irrelevant (purple, e.g., shape during colour task) stimulus dimensions, averaged over colour and shape. **B** MEG decoding of object colour information from the analogous empirical feature-based attention task of Goddard et al. (2022). Red shows decoding of the relevant (attended) stimulus (colour in the attend colour condition), and purple the irrelevant (unattended) stimulus (colour in the attend shape condition). Results show decoding accuracy for source estimated activity in a frontal ROI (inset). **C** Stimulus decoding accuracy from feature units, conventions as for A. **D** Same as B but for the estimated activity in a posterior ROI (inset). In B and D, lines in other colours refer to other attentional conditions not modelled here (pertaining to a distractor object), and grey shows estimated null distribution of decoding accuracies under permutation. Panels B and D are reproduced with permission from Goddard et al. (2022).

In summary, the minimal model was sufficient to reproduce the essential task-relevant prioritisation results that characterise human neuroimaging. In conjunction units, this manifested as differential activation for similar input depending on task rules, coding equivalent information more strongly when it was relevant for the task. In other words, the model gave rise to adaptive coding (Duncan 2001), as proposed for frontoparietal control of attention. Adaptive coding in the conjunctive layer in turn provided preferential excitatory input to the task relevant feature unit. That is, local competition in the feature units benefited from top down bias, as predicted by the biased competition model of attention (Desimone and Duncan 1995). This promoted a mutually virtuous cycle sustaining preferential representation of attended information (selective attention) throughout the system.

### Causal perturbation of the model reproduces non-invasive neurostimulation in humans

Next, we carried out causal test of the model by perturbing activation in the conjunction units. We compared the perturbation of the model to a unique human concurrent TMS-fMRI dataset (Jackson et al. 2021), which had investigated the causal role of right dlPFC in the prioritisation of relevant over irrelevant information in the rule-based selective attention task modelled above. In the human data, participants were provided with a rule indicating which stimulus dimension – colour or shape – they should respond to, and on each trial participants made a perceptual decision about the stimulus dimension indicated by the rule. While they were performing this task, participants received a train of three TMS pulses to right dlPFC at 13Hz, from 75ms after stimulus presentation at either a high (disruptive) or low (control) TMS intensity. Using multivariate decoding, the authors reported that in the multiple demand network, TMS primarily modulated decoding of the relevent information, rather than irrelevant information, suggesting that right dlPFC contributes specifically to supporting processing of relevant information (as opposed to suppressing irrelevant information). Behaviourally, the authors had expected an increase in congruency effect with TMS, but, observed that, if anything, the difference in RTs between congruent and incongruent trials was decreased, with congruent trials in particular becoming slower. Here, we asked whether perturbing the model would reproduce these effects, and if so, whether we could leverage the simplicity and interpretability of the model to provide a mechanistic hypothesis for how these effects could arise in vivo.

In our model, we simulated the effect of TMS by maximally activating the conjunction units in the stimulus presentation period. The maximal activation led to a period of suppression (arising from the blanket lateral inhibition between these units), the duration of which was determined by the TMS dose (length of the maximal activation, see Methods).

### TMS selectively disrupts coding of relevant information

First, we asked what effect perturbation of the model would have on the information that we could decode from unit activity. Inspection of model activity (**Figure 4)** showed that with moderate levels of TMS (**Figure 4A**), after maximal activation and the resulting suppression, conjunction units recovered such that the model could still solve the task. However, TMS appeared to flatten the difference between feature unit activations relative to the simulation without TMS (**Figure 2**). At high TMS levels (**Figure 4B**), conjunction to feature connectivity appeared to be so correlated that the model appeared to struggle to form attractors during the rule presentation period and was unable to perform the task.

**Figure 4.**
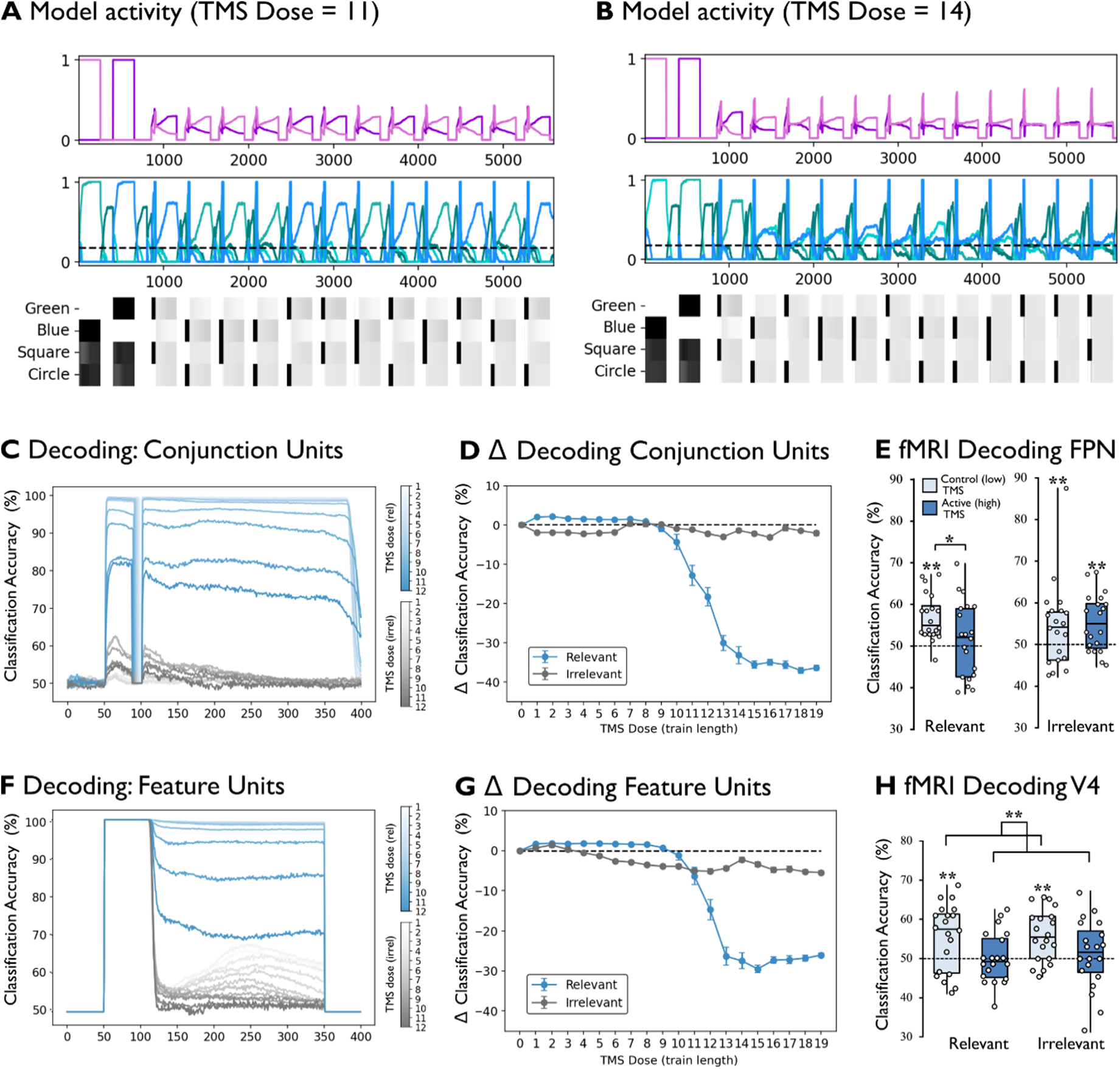
Effect of TMS on model activity and information coding. **A.** Model activity with a TMS dose of 11 timesteps, conventions as in Figure 2. **B.** Model activity with a TMS dose of 14 timesteps, conventions as in Figure 2. **C.** Decoding accuracy from conjunction units for relevant information (blue) and irrelevant information (dark grey). The shade of the colour corresponds to the value of TMS dose (colour bars). To avoid visual clutter and aid in interpretability only the first 12 values are shown. **D.** Average change in decoding accuracy for conjunction units relative to TMS = 0 averaged over the whole trial period for relevant information (blue) and irrelevant information (dark grey). **E** fMRI decoding accuracy in FPN regions, reproduced from Jackson et al. (2021). Light blue denotes the control condition and dark blue the active TMS condition. Significance markers above single bars denote decoding significantly above chance, and significance markers above brackets denote significance in post-hoc paired t-tests. Active TMS results in a reduction of relevant but not irrelevant information coding. **F., G.** same as C., D., but for the model’s sensory Feature Units. **H.** same as E but for visual region V4. * Denotes p<.05, and ** p<.008. rel: relevant; irrel: irrelevant.

To quantify these effects in terms of information coding, we separated coding of relevant and irrelevant stimulus information, in conjunctive and feature units, as we had done before, but now additionally we asked how coding was affected when TMS was applied at different doses. For conjunction units (**Figure 4 C,D)** low doses of TMS produced relatively small effects: the network was overall robust to low levels of perturbation. However, as TMS dose increased, particularly for doses of 10 or more, decoding of the relevant information in the conjunction units dropped sharply (**Figure 4 D**, blue line). In these units, coding of irrelevant information was relatively unaffected (**Figure 4 D**, grey line). In the feature units (**Figure 4 F,G)** low doses of TMS produced small effects. Moderate levels of TMS caused a drop in both relevant and irrelevant information coding (**Figure 4 G**, blue and grey lines). High levels of TMS again caused a sharp drop in relevant information coding in these regions. This pattern, particularly for TMS values of around 11, mimicked the human TMS-fMRI results reported by Jackson et al (2021) for FPN (**Figure 4 E**) and visual cortex (**Figure 4 H**).

### TMS affects model response times and accuracy

RTs increased with increasing levels of TMS until TMS dose 11 when they suddenly fell as accuracy dropped (**Figure 5A**). Most intriguingly, we also observed that in the model the size of the congruency effect for RT decreased under TMS (**Figure 5A**), similar to the counterintuitive empirical behavioural data from Jackson et al (2021) (**Figure 5B**). For low TMS doses, the accuracy of the model was relatively unaffected (**Figure 5C**), but accuracy rapidly dropped at doses greater than 10, falling to only marginally above chance for doses of 16 and higher.

**Figure 5.**
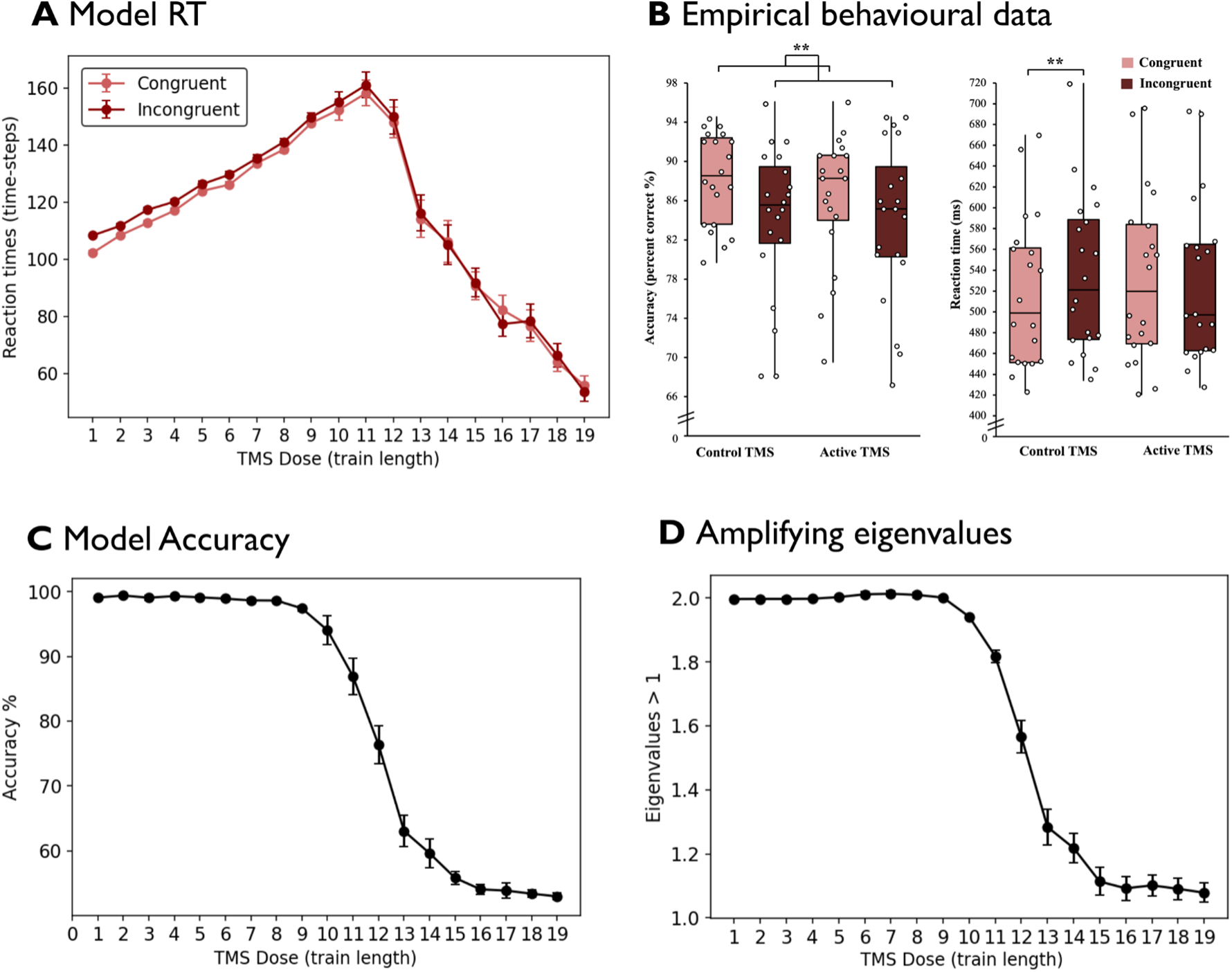
Effect of TMS on behaviour. **A** RTs for congruent and incongruent conditions at each TMS dose. **B** Empirical accuracy and RT data from Jackson et al, (2021), depicting human performance on congruent and incongruent trials in active TMS and control conditions. ** Denotes p<.001 in a paired t-test. **C** Model accuracy at each TMS dose. **D** Average number of eigenvalues >1 at the end of each block of trials for each TMS dose.

### Synaptic changes underpin TMS effects on both activity and behaviour

Finally, we leveraged the simplicity and interpretability of the model to generate a hypothesis for how these effects arose. We reasoned that maximal activation of the conjunctive units (TMS) would have two effects. First, there was a direct effect on activation in the current trial. In particular, lateral inhibition between conjunction units meant that once TMS was removed, their activity dropped to zero forcing them to re-compete (**Figure 4A, B)**. This delayed the onset of feedback to motor units, lengthening model reaction times. However, in parallel, we observed second – synaptic - mechanism by which TMS affected the model. In particular, based on the rapid Hebbian plasticity, we reasoned that the maximal coactivation of the conjunction units generated by the TMS pulse strengthened the connections between all conjunction units and the active feature units reducing the targeted connections that had been imprinted on the weight matrix by the rule presentation at the beginning of each block. This would weaken (i.e. make more shallow), and eventually destroy, the attractors underlying task performance. Accordingly, a flattening of the synaptic weights (i.e. a uniform increase in all synapses) was evident through visual inspection of the weights on each trial. To quantify this, we reduced the networks to a simple linear dynamical system with attractors that can be described by the eigenvalues of the weight matrix (which describes the asymptotic behaviour of the system). Specifically, we approximated the model with a reduced set of coupled difference equations (see Methods), and recorded the number of eigenvalues of the matrix that were > 1 at the end of each block of trials. This value quantifies the integrity of the attractors in the model. Strikingly, the average number of amplifying eigenvalues (**Figure 5D**) followed an almost identical pattern to task accuracy (**Figure 5C**) and relevant information coding (**Figure 4D, G**, blue lines) across the range of TMS doses. The behaviour of the network under TMS, therefore, has a straightforward interpretation. Namely, the coincident activation of the conjunction units increased the strength of the connections between the active feature units and all the conjunction units, reducing the specificity of the weight matrix. The network was only able to complete the task whilst the weight matrix underlying the recurrent circuit had as least two amplifying eigenvalues corresponding to the two attractors in the network’s dynamics. At low doses of TMS (<14), the model was able to recover and fall back into one of the attracting states. At TMS doses of 14 or more, the average number of amplifying eigenvalues was barely above 1, and consequently, accuracy approached chance. Under these conditions the network RT decreased because the motor units had a lower maximum value so it took less time to reach 98% of the maximum value. Reduction of the targeted connections in the weight matrix (weakening of the attractors) also provides a possible explanation for the observed reduction in the congruency effect. Specifically, the reduction in specificity of feedback from conjunction to feature neurons, or more precisely global increase in the strength of the synapses connecting the conjunction units with all of the features in the network, meant that after the removal of the stimulus, and recovery of activity in the winning conjunction unit, feedback from this conjunction unit to the relevant feature unit was less precise: feedback tended to excite all the feature units more diffusely. Because of this, as TMS dose increased, the difference in activity between the active feature unit and the non-active feature unit decreased. This diffuse feedback diminished any previous advantage provided by congruent relevant and irrelevant information or disadvantage for incongruent relevant and irrelevant information. The effect of these synaptic changes can be observed as “carry-over” effects in the model activity, with decoding on each trial being affected at timesteps during stimulus presentation before the TMS is presented on that trial (**Figure 4C**).

To further probe the role of synaptic plasticity, we ran an extra simulation in which we held the weight matrix constant (i.e., removed plasticity) after two rule presentation epochs in each block. Under these circumstances, of the model could still perform the task (mean accuracy 95%), but the model now failed to reproduce the empirical observation of a decrease in congruency effect with TMS. In fact, we observed that now congruency effects tended to increase with increasing TMS dose (**Figure S4A**). This observation adds weight to the argument that trial-to-trial plasticity in the model is necessary to account for the pattern of human data.

The change in relevant information coding in the network (classification accuracy) followed an almost identical pattern of results to behavioural accuracy. Specifically, relevant classification accuracy was maintained until the TMS dose reached a value of 10, after which which point it rapidly fell, until it was only slightly above chance. The explanation for this is almost identical to the explanation for the reduction in the congruency effect. Conjunction units developed stronger but less specific connections to all the feature units leading to a reduction in the prioritisation of relevant information as a function of TMS dose and reduced dramatically at the point at which the network was no longer able to maintain two attractors. In the absence of plasticity, the effect of TMS on decoding in conjunction and feature units was greatly diminished (**Figure S4B, C**) as attractors were maintained in the fixed weight matrix.

More speculatively, we can look for a location in the parameter space that reproduces the empirical results of Jackson et al. (2021). That is, where 1) classification accuracy for conjunction units is reduced but irrelevant information is relatively unperturbed, 2) classification accuracy in feature units is reduced for both relevant and irrelevant information, and 3) the congruency effect in RTs is reduced and absent. This set of constraints is broadly satisfied at a TMS dose of 10 or 11, which is also the point that, empirically, the average number of amplifying eigenvalues began to reduce. The only mismatch with the empirical data is that in the model TMS increased the overall length of RTs, but in the empirical data the overall length of RTs was not changed by TMS. We attribute this to the fact that the empirical task forced participants to make speeded responses within 500ms (Jackson et al. 2021) which was not implemented in the model.

## Discussion

The human capacity for flexible cognitive control - our ability rapidly prioritise different aspects of the world under different circumstances and associate them with different meaning and action – was traditionally regarded as a rather mysterious property of the human brain and cognition. Here, we show that it can arise very simply, in a tiny neural network imbued with three biologically plausible features: extensive connectivity between specialised “features” and associative “conjunctive” units, lateral inhibition, and rapid synaptic changes. These features are sufficient to give rise to selective attention via adaptive coding (in conjunction units) and biased competition (between feature units), two cornerstone concepts of attentional theory. By harnessing rapid synaptic plasticity to achieve zero-shot performance, this network accurately predicted human behavioural effects, neural decoding and their perturbation by TMS. The simplicity of the model allowed us to identify the precise mechanisms that generated these effects, and suggests that rapid synaptic changes provide a simple biologically plausible mechanism underpinning the control of flexible attention.

Our work establishes the face validity of the simple network as a model of rule based selective attention. In line with a wealth of human neuroimaging and non-human primate data (e.g., Moran and Desimone 1985, Chelazzi et al. 1993, Desimone and Duncan 1995, Luck et al. 1997, Chelazzi et al. 1998, Jehee et al. 2011, Woolgar et al. 2015, Jackson et al. 2017, Jackson and Woolgar 2018, Chen et al. 2021, Moerel et al. 2024, Zheng et al. 2024, Dermody et al. 2025), task-relevant information was prioritised in conjunction and feature units at the level of both firing rates and classification accuracy. The key processes underlying this effect were, 1) the initial broad-selectivity of the conjunction units which allowed the binding of stimuli and responses, 2) the Hebbian learning rule which created targeted connections between conjunction units and task-relevant features, and diffuse connections to task-irrelevant features, and 3) local competition between feature units. The targeted connectivity of task-relevant features generated a high and sustained firing rate response in the relevant dimension, and the diffuse connectivity to irrelevant features reduced the firing rate response in the irrelevant dimension as both stimulus features received similar feedback from the active conjunction unit and so inhibited each other relatively equally.

Notably, flexible selective attention arose in the model without any explicit representation of how attention should be applied (e.g., no “attention unit”), or indeed any explicit sustained representation of the current rule in the firing rates. Instead, the key manipulation responsible for this context sensitive behaviour was the presentation of the rules at the beginning of each block. The maximal activation of the relevant stimulus-response mapping, paired with the partial coactivation of the irrelevant dimension, resulted in targeted excitatory synaptic connections between the relevant stimulus and the winning conjunction unit, and diffuse synaptic connections to the irrelevant dimension. The connectivity change, combined with the local competition present in the feature units, meant that when each stimulus was presented, the stimulus feature in the relevant dimension received targeted “top-down” excitatory feedback from the active conjunction unit, whilst the non-relevant dimension received diffuse feedback. This led to a sustained high firing response in the relevant dimension even after the stimulus was removed. In contrast, diffuse feedback to the irrelevant dimension meant that there was no particular support for representation of the recently presented feature, which in turn did not reliably win the race to dominance through lateral inhibition. Thus in the model, flexible selective attention arises as a consequence of momentary stimulus-response associations arising from recently having performed (or preparing to perform) those associations.

The model formalises key proposals in the cognitive control and selective attention literature. These theories (Duncan 2010, Duncan et al. 2020, Cole 2024, Duncan 2024) converge on the concept that a set of broadly distributed and strongly and flexibly connected frontoparietal brain regions (variously called the multiple demand network (Duncan 2010), cognitive control network/networks (Cole and Schneider 2007, Gratton et al. 2018), frontoparietal control network (Vincent et al. 2008)) integrates task relevant elements into the structure required for the current task. When participants are given task instructions, the multiple demand regions are hypothesised to assemble the relevant task components into a “mental program” which then guides the completion of the relevant sub-tasks such as to what to attend to and what actions to execute (Duncan 2010, Duncan et al. 2020, Duncan 2024). Here we formalised this conceptual model in terms of a plastic attractor. In contrast to the original hypothesis, which was cast in terms of analogies to symbolic approaches in artificial intelligence (e.g., Newell 1994), here the formation of adaptive codes, and their role in task completion, is cashed out in dynamic terms. When the network is shown a set of rules the relevant sensory features and responses are bound together through Hebbian learning to a shared conjunction unit forming an attractor in the network’s dynamics. Stimulus presentation then excites the relevant conjunction neuron which, through excitatory feedback to the relevant feature neurons, pulls the network into the relevant attractor or “mental program” driving task completion.

This formalisation gives rise to two long-posited concepts in attentional theory: adaptive coding (Duncan 2001), and biased competition (Desimone and Duncan 1995). Adaptive coding is the proposal that single neurons in prefrontal (and other control) regions adapt their responses to prioritise discrimination of the currently relevant information. In our current model, this manifests as flexible selectivity profiles in the conjunction units, which reflect information in the attended dimension. Similar inputs lead to different activation in conjunction units according to the current task rules, based on the model’s current connectivity profile (i.e., the differential attractor states created by recently using or preparing to use the task rules). Biased competition is the proposal that local competition (e.g., in visual cortices) can be biased from top-down (e.g., prefrontal) influences in favour of the attended information (Desimone and Duncan 1995, Kastner et al. 1998, Reynolds et al. 1999). In our model, the two dimensions of a stimulus (its colour and shape) do not compete directly. However, the relevant information (e.g., colour) is supported by targeted excitation of the relevant presented feature unit (e.g., green), while competition between irrelevant features (e.g., shape) receives weak and diffuse input that does not support the presented feature (e.g., square). Thus, only competition in the relevant dimension is biased and therefore representation of the attended task feature is favoured. Rather than a strictly “top-down” or “bottom-up” effect, this creates a mutually virtuous cycle between activation in the feature unit in the relevant dimension, the relevant conjunction unit and the response unit, and the connections between them.

By perturbing the model with TMS we could further understand the simultaneous TMS-fMRI findings of Jackson et al. (2021). The empirical study was designed to test the role of dlPFC in the prioritisation of task-relevant information in the multiple demand network, and its role in guiding selective attention in sensory cortices. In terms of the effects relevant to the empirical study, the model showed a robust general pattern of reduced classification accuracy for task-relevant information in both feature and conjunction units, and a reduction in the size of the congruency effect over a wide range of the TMS doses. The full constellation of effects was, however, less robust and was only reproduced by a narrow range of the TMS doses. Assuming that the architecture of the model is broadly representative of the cognitive architecture of the human brain, these simulations point to the effects that we should expect to be replicable and those that are likely to be fragile and sensitive to small deviations in parameters. The crucial contribution of the model here, however, is in the explanation of the results that emerge. Specifically, the reduction in decoding accuracy for relevant information, and the reduction in the size of the congruency effect, are both due to TMS globally strengthening the synapses between all the conjunction and feature units, reducing the specificity of their connectivity. This gives rise to the specific hypothesis that synaptic connectivity may underpin control of attention.

A number of limitations remain. In modelling the phenomenological effects of TMS, even if we set aside questions about biophysical properties (e.g. the role of spike timing, and refractory periods), the current model made a number of stark idealisations. In reality, the TMS induced electric field does not perturb all neurons equally, as modelled here. The strength of the induced electric field is strongest at the central site of stimulation and weaker elsewhere (e.g., Opitz et al. 2011). However, the effects are far from uniform and neural responses depend on many anatomical factors including the orientation of the neurons relative to the magnetic field (e.g, Weise et al. 2020, Turi et al. 2022, Dannhauer et al. 2024). Also relevant is the fact that in the experimental paradigm of Jackson et al. (2021) TMS was applied to right dlPFC, and the reduction of decoding accuracy for relevant information was observed throughout the rest of the multiple demand network (although similar results were found also in the directly targeted region). This, again, was not modelled in the current work. It is also possible that our reliance on purely Hebbian plasticity leads to an unrealistic level of correlated activity in the network under long TMS doses.

All models have relative advantages and disadvantages. Previous simulations have created neural networks that switch between tasks over a single trial (e.g., Song et al. 2016, Mohan et al. 2021), but most have required extensive training on the exact tasks. Networks have also been created that generalise to novel tasks by recombining known features (e.g., Oh et al. 2017, Lake and Baroni 2018) but these have tended to be large and complex, and may not have biologically plausible learning algorithms. Our model is simple, does not require extensive training and relies on biologically plausible Hebbian learning. One strength of the model is its ability to act using novel, never-before-seen stimulus-response mappings. However, since our model is highly simplified, it requires modification to handle larger sets of tasks, unlike deep networks or larger unstructured RNNs. For example, our model lacks abstraction, and works only on features that are already represented within the network. As such, in its current form, it would be unable to solve exclusive-or (XOR) tasks without ‘flattening’ the task representation. The model would also require modification in order to switch between task sets based on being presented with a rule cue rather than by experiencing the entire stimulus-response mapping, although this could conceivably be achieved via a minor modification of adding an additional feature unit that particular cues activate. Further work will be needed to test the limits of what aspects of flexible control the principles underlying the operation of the model can give rise to, including whether the model can generalise to multi-response tasks, where the response domain is more complex, and where the stimulus set is more complex. Due to our model being rate-coded, it also cannot provide predictions about neural synchrony or oscillations, unlike other models (Deco and Rolls 2003, Wei et al. 2012, Bays 2015, Fiebig and Lansner 2017). On the other hand, our model is simple enough for each neuron’s role to be understood. It allows simple testing of predictions about TMS and decoding, as well as behaviour. Importantly, the mechanistic operation of the network is analysable using well-established methods of dynamical system theory.

In sum, we found that modelling rapid synaptic changes leads to a powerful formalism to model rule based selective-attention. Despite its simplicity, the model provides concrete hypotheses about the role the conjunctive codes in prioritising relevant information and accurately predicted a range of neural findings in a simple and explainable way, contributing a plausible mechanism underlying flexible selective attention.

## Acknowledgements

This work was funded by UKRI MRC intramural funding (SUAG/093/G116768 to A.W.). For the purpose of open access, the author has applied a Creative Commons Attribution (CC BY) licence to any Author Accepted Manuscript version arising from this submission.

## Supplementary material

### Long(er) Time-scale Plasticity is required for behavioural congruency effects and consistent conjunction unit selectivity across blocks

We tested the behaviour of an unmodified plastic attractor - based on the equations and parameters supplied in the supplementary material of Manohar et al. (2019) – on the feature-based attention task introduced in the main section of the paper. In this network the synapses connecting feature and conjunction units change on a single, relatively fast, time-scale. The unmodified model is able to perform the task with a high degree of accuracy (M = 97.7%, SEM = 0.002) However, without the addition of a long(er) timescale of plasticity that provides a bias to the competition between conjunction units, encouraging their reuse across blocks, the model does not show a congruency effect, and the selectivity of conjunction units changes overtime (**Figure S1)**. Both of these aspects of the network’s behaviour contradict established empirical findings, driving us to modify the original model. In particular, behavioural congruency effects are well established, including in the Jackson et al. (2021) task we aimed to reproduce through simulation, and at the level of large neuronal populations, selectivity is somewhat stable over time as evidenced by fMRI decoding (including from the multiple demand regions) which is typically cross-validated across runs of several minutes.

**Figure S1.**
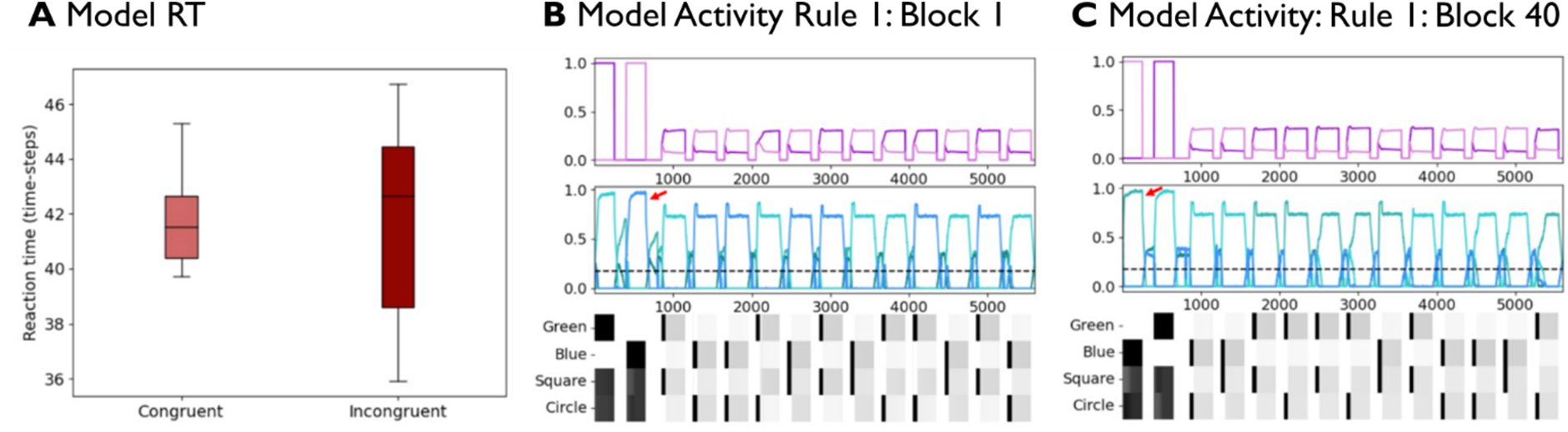
RT and model activity with faster weight updating. **A.** Boxplot summary of simulated RTs (number of time-steps to 98% of maximum activation) for congruent and incongruent trials. **B.** Example model activity on block 1 of the feature based attention task. Bottom raster style plots show the activity of the feature units, the middle line plots show conjunction unit activity, and the top line plots show motor response activity. Each colour represents a distinct conjunction or motor unit. **C.** Model activity on block 40 of the same task, conventions as in B. Notice that the colour of the dominant conjunction units has changed (red arrows) despite the stimulus features, responses, and rules remaining consistent signifying a change in the selectivity of conjunction units. By trial 40, the dominant conjunction unit paired with the pink motor response changed from the light blue conjunction unit to the dark teal conjunction unit.

### Modified model is still able to perform original working memory task

We tested whether our modified model was still able to perform the working memory task used by Manohar et al. (2019) in the original presentation. On each trial, the network was presented with four objects each consisting of 2 features. After the presentation there was a brief delay and a probe, consisting of one feature of the four objects, was presented. The task was then to recall the full object. The network completed the task through pattern completion. The presentation of the probe excited the conjunction unit that binds together all of the features of the object bringing the other feature of the object above its baseline firing rate. Successful completion of the task was defined as the reactivation of the feature originally paired with the probe feature as is shown below. As is shown in **Figure S2**, even with the addition of long(er) term plasticity in the modified model, the conjunction units still had the requisite flexibility to perform the working memory task that originally motivated the model.

**Figure S2.**
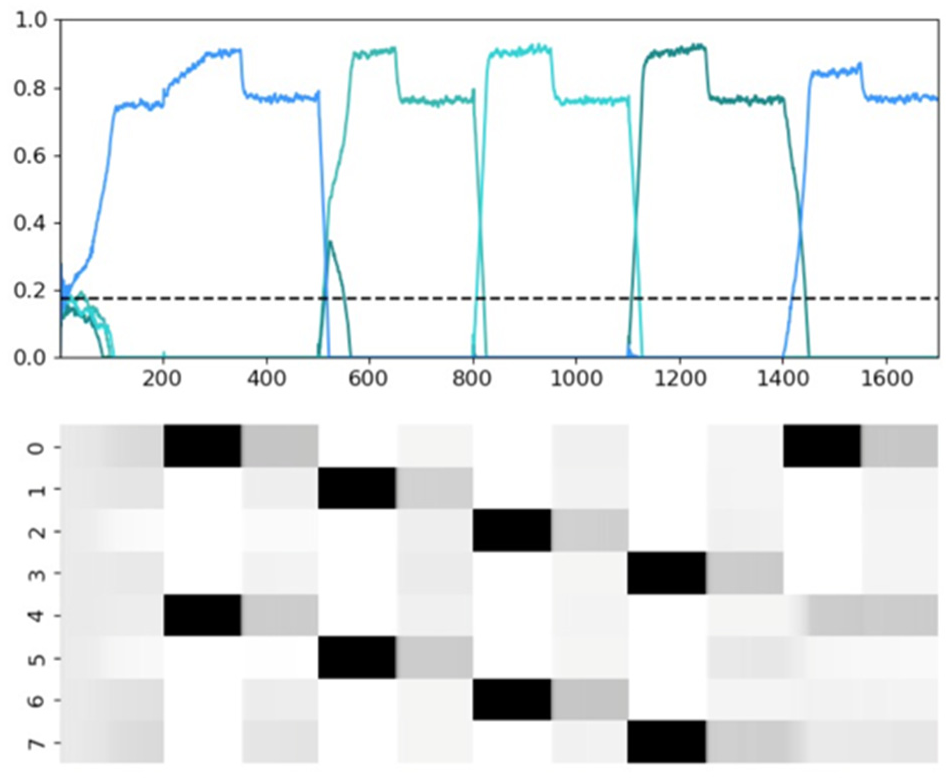
Model activity for the original working memory task. Simplified working memory task adapted from Manohar et al (2019), implemented with the modified model used in this paper. Rasta style plots show the firing rates of feature units, and the line plots above show the behaviour of the conjunction units with each colour corresponding to a distinct conjunction unit. The modified model is still able to perform this task.

### Interleaving TMS trial-wise

We tested whether a similar pattern of results would be obtained if TMS was presented on only a subset of trials, as is more common in human TMS experiments. The behaviour of the model on TMS trials was similar; in particular, it reproduced the change in congruency effect.

**Figure S3.**
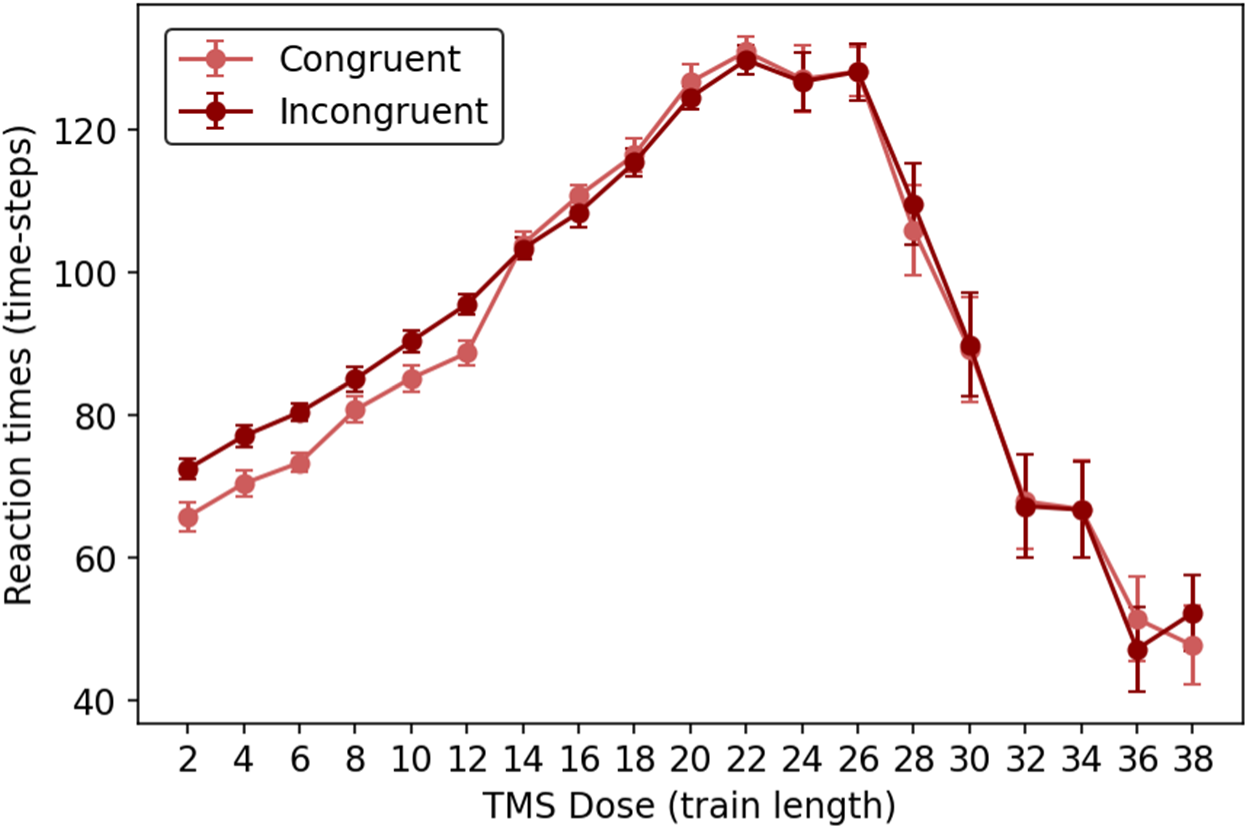
Effect of interleaved TMS on RT. RTs for congruent and incongruent conditions at each TMS dose, when TMS is only presented on 50% of trials (interleaved). The model reproduced a reduction in congruency effect at higher TMS doses.

### Trial-to-trial plasticity is necessary to capture the effects of TMS on decoding and behaviour

We tested whether a similar pattern of results could be obtained without plastic updating of the weight matrix for any trials or timepoints after rule presentation. This model could still perform the task (mean accuracy 95%). However, instead of decreasing with TMS, as in the human behavioural data and with the main model, the congruency effect tended to *increase* with increasing TMS dose in the model without plasticity (**Figure S4A**). The effect of TMS on information coding decoding was markedly reduced, even at high TMS doses, in both conjunction (**Figure S4B**), and feature units (**Figure S4C**).

**Figure S4.**
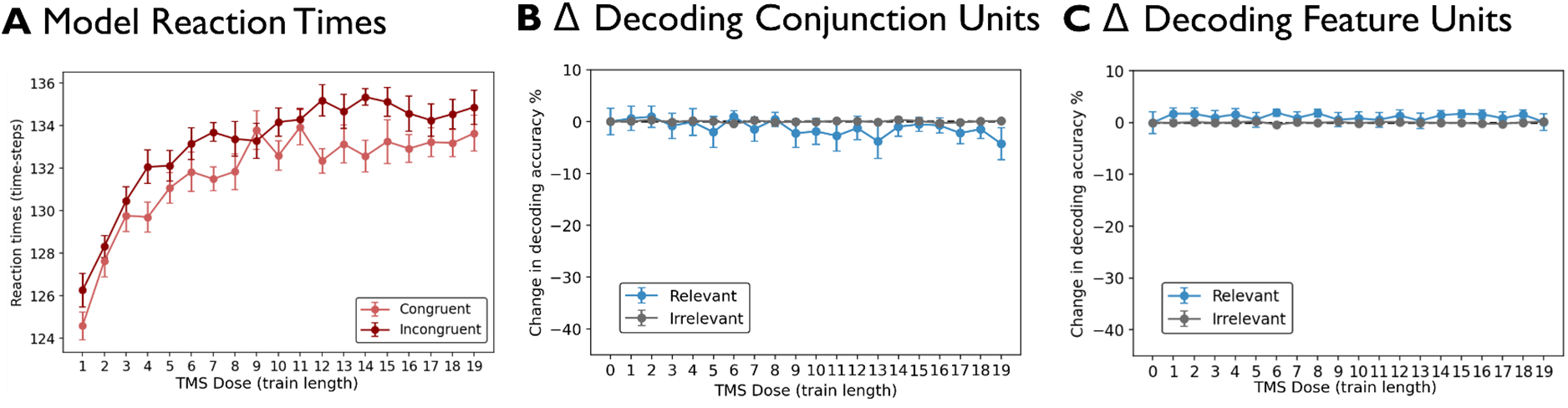
Effect of TMS on Model RT and decoding in the absence of plasticity. **A.** RTs for congruent and incongruent conditions at each TMS dose. Higher doses of TMS increase, rather than decrease, the behavioural congruency effect. **B.** Average change in decoding accuracy for conjunction units relative to no TMS (TMS = 0) averaged over the whole trial period for relevant information (blue) and irrelevant information (dark grey). **C.** Same as B but for feature units.

## References

Altmann, C. F., M. Henning, M. K. Döring and J. Kaiser (2008). “Effects of feature-selective attention on auditory pattern and location processing.” NeuroImage 41(1): 69–79.

Anllo-Vento, L. and S. A. Hillyard (1996). “Selective attention to the color and direction of moving stimuli: Electrophysiological correlates of hierarchical feature selection.” Perception & Psychophysics 58(2): 191–206.

Assem, M., M. F. Glasser, D. C. Van Essen and J. Duncan (2020). “A domain-general cognitive core defined in multimodally parcellated human cortex.” Cerebral Cortex 30(8): 4361–4380.

Badre, D. and M. D’Esposito (2007). “Functional magnetic resonance imaging evidence for a hierarchical organization of the prefrontal cortex.” J Cogn Neurosci 19(12): 2082–2099.

Barnes, L., E. Goddard and A. Woolgar (2021). “Neural coding of visual objects rapidly reconfigures to reflect sub-trial shifts in attentional focus.” BioRxiv.

Battistoni, E., D. Kaiser, C. Hickey and M. V. Peelen (2020). “The time course of spatial attention during naturalistic visual search.” Cortex 122: 225–234.

Bays, P. M. (2015). “Spikes not slots: noise in neural populations limits working memory.” Trends in Cognitive Sciences 19(8): 431–438.

Bergmann, T. O., R. Varatheeswaran, C. A. Hanlon, K. H. Madsen, A. Thielscher and H. R. Siebner (2021). “Concurrent TMS-fMRI for causal network perturbation and proof of target engagement.” Neuroimage 237: 118093.

Bocincova, A., T. J. Buschman, M. G. Stokes and S. G. Manohar (2022). “Neural signature of flexible coding in prefrontal cortex.” Proc Natl Acad Sci U S A 119(40): e2200400119.

Bode, S. and J. D. Haynes (2009). “Decoding sequential stages of task preparation in the human brain.” Neuroimage 45(2): 606–613.

Botvinick, M. M. and J. D. Cohen (2014). “The computational and neural basis of cognitive control: charted territory and new frontiers.” Cogn Sci 38(6): 1249–1285.

Bunge, S. A., I. Kahn, J. D. Wallis, E. K. Miller and A. D. Wagner (2003). “Neural circuits subserving the retrieval and maintenance of abstract rules.” J Neurophysiol 90(5): 3419–3428.

Chelazzi, L., J. Duncan, E. K. Miller and R. Desimone (1998). “Responses of neurons in inferior temporal cortex during memory-guided visual search.” J Neurophysiol 80(6): 2918–2940.

Chelazzi, L., E. K. Miller, J. Duncan and R. Desimone (1993). “A neural basis for visual search in inferior temporal cortex.” Nature 363(6427): 345–347.

Chen, J., P. S. Scotti, E. W. Dowd and J. D. Golomb (2021). “Neural Representations of Task-relevant and Task-irrelevant Features of Attended Objects.” bioRxiv.

Chen, X., K.-P. Hoffmann, T. D. Albright and A. Thiele (2012). “Effect of feature-selective attention on neuronal responses in macaque area MT.” Journal of Neurophysiology 107(5): 1530–1543.

Cohen, J. D., K. Dunbar and J. L. Mcclelland (1990). “On the Control of Automatic Processes - a Parallel Distributed-Processing Account of the Stroop Effect.” Psychological Review 97(3): 332–361.

Cole, M. W. (2024). “Cognitive flexibility as the shifting of brain network flows by flexible neural representations.” Current Opinion in Behavioral Sciences 57: 101384.

Cole, M. W., P. Laurent and A. Stocco (2013). “Rapid instructed task learning: A new window into the human brain’s unique capacity for flexible cognitive control.” Cognitive, Affective, & Behavioral Neuroscience 13(1): 1–22.

Cole, M. W., J. R. Reynolds, J. D. Power, G. Repovs, A. Anticevic and T. S. Braver (2013). “Multi-task connectivity reveals flexible hubs for adaptive task control.” Nature neuroscience 16(9): 1348–1355.

Cole, M. W. and W. Schneider (2007). “The cognitive control network: Integrated cortical regions with dissociable functions.” Neuroimage 37(1): 343–360.

Dannhauer, M., L. J. Gomez, P. L. Robins, D. Wang, N. I. Hasan, A. Thielscher, H. R. Siebner, Y. Fan and Z.-D. Deng (2024). “Electric Field Modeling in Personalizing Transcranial Magnetic Stimulation Interventions.” Biological Psychiatry 95(6): 494–501.

Deco, G. and E. T. Rolls (2003). “Attention and working memory: a dynamical model of neuronal activity in the prefrontal cortex.” European Journal of Neuroscience 18(8): 2374–2390.

Dehaene, S. and J. P. Changeux (1991). “The Wisconsin Card Sorting Test: theoretical analysis and modeling in a neuronal network.” Cereb Cortex 1(1): 62–79.

Denison, R. N., M. Carrasco and D. J. Heeger (2021). “A dynamic normalization model of temporal attention.” Nature Human Behaviour 5(12): 1674–1685.

Dermody, N., R. Lorenz, E. Goddard, A. Villringer and A. Woolgar (2025). “Spatial and feature-selective attention interact to drive selective coding in frontoparietal cortex.” Neuropsychologia 216: 109172.

Desimone, R. and J. Duncan (1995). “Neural mechanisms of selective visual attention.” Annu Rev Neurosci 18: 193–222.

Downer, J. D., B. Rapone, J. Verhein, K. N. O’Connor and M. L. Sutter (2017). “Feature-Selective Attention Adaptively Shifts Noise Correlations in Primary Auditory Cortex.” The Journal of Neuroscience 37(21): 5378–5392.

Duncan, J. (2001). “An adaptive coding model of neural function in prefrontal cortex.” Nat Rev Neurosci 2(11): 820–829.

Duncan, J. (2010). “The multiple-demand (MD) system of the primate brain: mental programs for intelligent behaviour.” Trends Cogn Sci 14(4): 172–179.

Duncan, J. (2024). “Construction and use of mental models: Organizing principles for the science of brain and mind.” Neuropsychologia: 109062.

Duncan, J., M. Assem and S. Shashidhara (2020). “Integrated Intelligence from Distributed Brain Activity.” Trends Cogn Sci 24(10): 838–852.

Fiebig, F. and A. Lansner (2017). “A Spiking Working Memory Model Based on Hebbian Short-Term Potentiation.” The Journal of Neuroscience 37(1): 83–96.

Fiebig, F. and A. Lansner (2017). “A spiking working memory model based on hebbian short-term potentiation.” Journal of Neuroscience 37(1): 83–96.

Flesch, T., K. Juechems, T. Dumbalska, A. Saxe and C. Summerfield (2022). “Orthogonal representations for robust context-dependent task performance in brains and neural networks.” Neuron 110(7): 1258–1270.e1211.

Freedman, D. J., M. Riesenhuber, T. Poggio and E. K. Miller (2001). “Categorical representation of visual stimuli in the primate prefrontal cortex.” Science 291(5502): 312–316.

Garner, W. R. (1978). “Selective attention to attributes and to stimuli.” Journal of Experimental Psychology: General 107(3): 287.

Goddard, E., T. A. Carlson and A. Woolgar (2022). “Spatial and Feature-selective Attention Have Distinct, Interacting Effects on Population-level Tuning.” J Cogn Neurosci 34(2): 290–312.

Gonzalez-Garcia, C., J. E. Arco, A. F. Palenciano, J. Ramirez and M. Ruz (2017). “Encoding, preparation and implementation of novel complex verbal instructions.” Neuroimage 148: 264–273.

Gonzalez-Garcia, C., S. Formica, B. Liefooghe and M. Brass (2020). “Attentional prioritization reconfigures novel instructions into action-oriented task sets.” Cognition 194: 104059.

Gonzalez-Garcia, C., S. Formica, D. Wisniewski and M. Brass (2021). “Frontoparietal action-oriented codes support novel instruction implementation.” Neuroimage 226: 117608.

Gratton, C., K. K. Sreenivasan, M. A. Silver and M. D’Esposito (2013). “Attention Selectively Modifies the Representation of Individual Faces in the Human Brain.” The Journal of Neuroscience 33(16): 6979–6989.

Gratton, C., H. Sun and S. E. Petersen (2018). “Control networks and hubs.” Psychophysiology 55(3): e13032.

Grootswagers, T., A. K. Robinson, S. M. Shatek and T. A. Carlson (2021). “The neural dynamics underlying prioritisation of task-relevant information.” Neurons, Behavior, Data analysis, and Theory 5(1): 1–17.

Grootswagers, T., S. G. Wardle and T. A. Carlson (2017). “Decoding Dynamic Brain Patterns from Evoked Responses: A Tutorial on Multivariate Pattern Analysis Applied to Time Series Neuroimaging Data.” J Cogn Neurosci 29(4): 677–697.

Howard, M. W. and M. J. Kahana (2002). “A Distributed Representation of Temporal Context.” Journal of Mathematical Psychology 46(3): 269–299.

Ito, T., G. R. Yang, P. Laurent, D. H. Schultz and M. W. Cole (2022). “Constructing neural network models from brain data reveals representational transformations linked to adaptive behavior.” Nat Commun 13(1): 673.

Jackson, J., E. Feredoes, A. N. Rich and A. Woolgar (2021). “Concurrent neuroimaging and neural stimulation reveals a causal role for dlPFC in selection of task-relevant information “ Nature Communications Biology 4: 588.

Jackson, J., A. N. Rich, M. A. Williams and A. Woolgar (2017). “Feature-selective Attention in Frontoparietal Cortex: Multivoxel Codes Adjust to Prioritize Task-relevant Information.” Journal of Cognitive Neuroscience 29(2): 310–321.

Jackson, J. and A. Woolgar (2018). “Adaptive coding in the human brain: Distinct object features are encoded by overlapping voxels in frontoparietal cortex.” Cortex 108: 25–34.

Jackson, J. B., E. Feredoes, A. N. Rich, M. Lindner and A. Woolgar (2021). “Concurrent neuroimaging and neurostimulation reveals a causal role for dlPFC in coding of task-relevant information.” Commun Biol 4(1): 588.

Jackson, J. B., A. N. Rich, D. Moerel, L. Teichmann, J. Duncan and A. Woolgar (2025). “Domain general frontoparietal regions show modality-dependent coding of auditory and visual rules.” Imaging Neuroscience 3.

Jehee, J. F., D. K. Brady and F. Tong (2011). “Attention improves encoding of task-relevant features in the human visual cortex.” J Neurosci 31(22): 8210–8219.

Kastner, S., P. De Weerd, R. Desimone and L. G. Ungerleider (1998). “Mechanisms of Directed Attention in the Human Extrastriate Cortex as Revealed by Functional MRI.” Science 282(5386): 108–111.

Kauramäki, J., I. P. Jääskeläinen and M. Sams (2007). “Selective Attention Increases Both Gain and Feature Selectivity of the Human Auditory Cortex.” PLOS ONE 2(9): e909.

Koechlin, E. and C. Summerfield (2007). “An information theoretical approach to prefrontal executive function.” Trends Cogn Sci 11(6): 229–235.

Lake, B. and M. Baroni (2018). Generalization without Systematicity: On the Compositional Skills of Sequence-to-Sequence Recurrent Networks. Proceedings of the 35th International Conference on Machine Learning. D. Jennifer and K. Andreas. Proceedings of Machine Learning Research, PMLR. 80: 2873--2882.

Lu, R., E. Michael, C. L. Scrivener, J. B. Jackson, J. Duncan and A. Woolgar (2025). “Parietal alpha stimulation causally enhances information coding in evoked and oscillatory activity “ Brain Stimulation.

Luck, S. J., L. Chelazzi, S. A. Hillyard and R. Desimone (1997). “Neural mechanisms of spatial selective attention in areas V1, V2, and V4 of macaque visual cortex.” J Neurophysiol 77(1): 24–42.

Manohar, S. G., N. Zokaei, S. J. Fallon, T. P. Vogels and M. Husain (2019). “Neural mechanisms of attending to items in working memory.” Neuroscience & Biobehavioral Reviews 101: 1–12.

Moerel, D., T. Grootswagers, A. K. Robinson, S. M. Shatek, A. Woolgar, T. A. Carlson and A. N. Rich (2022). “The time-course of feature-based attention effects dissociated from temporal expectation and target-related processes.” Scientific Reports 12(1): 6968.

Moerel, D., A. N. Rich and A. Woolgar (2024). “Selective Attention and Decision-Making Have Separable Neural Bases in Space and Time.” The Journal of Neuroscience 44(38): e0224242024.

Mohan, K., O. Zhu and D. J. Freedman (2021). “Interaction between neuronal encoding and population dynamics during categorization task switching in parietal cortex.” Neuron 109(4): 700–712.e704.

Moore, T. and K. M. Armstrong (2003). “Selective gating of visual signals by microstimulation of frontal cortex.” Nature 421(6921): 370–373.

Moran, J. and R. Desimone (1985). “Selective attention gates visual processing in the extrastriate cortex.” Science 229(4715): 782–784.

Newell, A. (1994). Unified theories of cognition, Harvard University Press.

Nobre, A. C., A. Rao and L. Chelazzi (2006). “Selective Attention to Specific Features within Objects: Behavioral and Electrophysiological Evidence.” Journal of Cognitive Neuroscience 18(4): 539–561.

O’Reilly, R. C. and M. J. Frank (2006). “Making Working Memory Work: A Computational Model of Learning in the Prefrontal Cortex and Basal Ganglia.” Neural Computation 18(2): 283–328.

Oh, J., S. Singh, H. Lee and P. Kohli (2017). Zero-Shot Task Generalization with Multi-Task Deep Reinforcement Learning. Proceedings of the 34th International Conference on Machine Learning. P. Doina and T. Yee Whye. Proceedings of Machine Learning Research, PMLR. 70: 2661--2670.

Opitz, A., M. Windhoff, R. M. Heidemann, R. Turner and A. Thielscher (2011). “How the brain tissue shapes the electric field induced by transcranial magnetic stimulation.” NeuroImage 58(3): 849–859.

Palenciano, A. F., C. Gonzalez-Garcia, J. E. Arco, L. Pessoa and M. Ruz (2019). “Representational Organization of Novel Task Sets during Proactive Encoding.” J Neurosci 39(42): 8386–8397.

Pedregosa, F., G. Varoquaux, A. Gramfort, V. Michel, B. Thirion, O. Grisel, M. Blondel, P. Prettenhofer, R. Weiss and V. Dubourg (2011). “Scikit-learn: Machine learning in Python.” the Journal of machine Learning research 12: 2825–2830.

Rao, S. C., G. Rainer and E. K. Miller (1997). “Integration of what and where in the primate prefrontal cortex.” Science 276(5313): 821–824.

Reynolds, J. H., L. Chelazzi and R. Desimone (1999). “Competitive Mechanisms Subserve Attention in Macaque Areas V2 and V4.” The Journal of Neuroscience 19(5): 1736–1753.

Romero, M. C., M. Davare, M. Armendariz and P. Janssen (2019). “Neural effects of transcranial magnetic stimulation at the single-cell level.” Nature Communications 10(1): 2642.

Rossi, A. F. and M. A. Paradiso (1995). “Feature-specific effects of selective visual attention.” Vision Research 35(5): 621–634.

Ruff, C. C., F. Blankenburg, O. Bjoertomt, S. Bestmann, E. Freeman, J. D. Haynes, G. Rees, O. Josephs, R. Deichmann and J. Driver (2006). “Concurrent TMS-fMRI and psychophysics reveal frontal influences on human retinotopic visual cortex.” Curr Biol 16(15): 1479–1488.

Sandberg, A., J. Tegnér and A. Lansner (2003). “A working memory model based on fast Hebbian learning.” Network: Computation in Neural Systems 14(4): 789.

Schultz, D. H., T. Ito and M. W. Cole (2022). “Global connectivity fingerprints predict the domain generality of multiple-demand regions.” Cerebral Cortex 32(20): 4464–4479.

Snyder, A. C., B. M. Yu and M. A. Smith (2021). “A Stable Population Code for Attention in Prefrontal Cortex Leads a Dynamic Attention Code in Visual Cortex.” The Journal of Neuroscience 41(44): 9163–9176.

Soltani, A. and E. Koechlin (2022). “Computational models of adaptive behavior and prefrontal cortex.” Neuropsychopharmacology 47(1): 58–71.

Song, H. F., G. R. Yang and X.-J. Wang (2016). “Training Excitatory-Inhibitory Recurrent Neural Networks for Cognitive Tasks: A Simple and Flexible Framework.” PLOS Computational Biology 12(2): e1004792.

Turi, Z., N. Hananeia, S. Shirinpour, A. Opitz, P. Jedlicka and A. Vlachos (2022). “Dosing Transcranial Magnetic Stimulation of the Primary Motor and Dorsolateral Prefrontal Cortices With Multi-Scale Modeling.” Frontiers in Neuroscience **Volume** 16-2022.

Veniero, D., J. Gross, S. Morand, F. Duecker, A. T. Sack and G. Thut (2021). “Top-down control of visual cortex by the frontal eye fields through oscillatory realignment.” Nature Communications 12(1): 1757.

Vincent, J. L., I. Kahn, A. Z. Snyder, M. E. Raichle and R. L. Buckner (2008). “Evidence for a frontoparietal control system revealed by intrinsic functional connectivity.” J Neurophysiol 100(6): 3328–3342.

Wallis, J. D. and E. K. Miller (2003). “From Rule to Response: Neuronal Processes in the Premotor and Prefrontal Cortex.” Journal of Neurophysiology 90(3): 1790–1806.

Wei, Z., X.-J. Wang and D.-H. Wang (2012). “From Distributed Resources to Limited Slots in Multiple-Item Working Memory: A Spiking Network Model with Normalization.” The Journal of Neuroscience 32(33): 11228–11240.

Weise, K., O. Numssen, A. Thielscher, G. Hartwigsen and T. R. Knösche (2020). “A novel approach to localize cortical TMS effects.” NeuroImage 209: 116486.

Wisniewski, D., C. Gonzalez-Garcia, S. Formica, A. Woolgar and M. Brass (2023). “Adaptive coding of stimulus information in human frontoparietal cortex during visual classification.” Neuroimage 274: 120150.

Woolgar, A., S. Afshar, M. A. Williams and A. N. Rich (2015). “Flexible Coding of Task Rules in Frontoparietal Cortex: An Adaptive System for Flexible Cognitive Control.” J Cogn Neurosci 27(10): 1895–1911.

Woolgar, A., E. Feredoes, M. Assem, Y. Bassil, T. O. Bergmann, L. Beynel, M. Burke, R. F. H. Cash, R. M. Comeau, M. M. Correia, E. Genc, G. Hartwigsen, J. B. Jackson, M. Kienle, P. Kunz, O. Leticevscaia, B. Luber, M. Lueckel, C. Mathiesen, E. Michael, O. Numssen, D. J. Oathes, A. C. Rosen, T. Schuhmann, A.-L. Schuler, C. L. Scrivener, A. Thielscher, M. Tik, Y. Todorov, M. Vasileiadi, C. Windischberger, M. S. Hermiller and A. T. Sack (2025). “Consensus guidelines for the use of concurrent TMS-fMRI in cognitive and applied human neuroscience “ Nature Protocols.

Woolgar, A., J. Jackson and J. Duncan (2016). “Coding of Visual, Auditory, Rule, and Response Information in the Brain: 10 Years of Multivoxel Pattern Analysis.” J Cogn Neurosci 28(10): 1433–1454.

Woolgar, A., M. A. Williams and A. N. Rich (2015). “Attention enhances multi-voxel representation of novel objects in frontal, parietal and visual cortices.” Neuroimage 109: 429–437.

Zheng, Y., R. Lu and A. Woolgar (2024). “Radical flexibility of neural representation in frontoparietal cortex and the challenge of linking it to behaviour.” Current Opinion in Behavioral Sciences 57: 101392.

